# Duration of immunity following infection with moderately virulent ASFV

**DOI:** 10.1101/2025.06.05.657993

**Authors:** Virginia Friedrichs, Paul Deutschmann, Kerstin Wernike, Tessa Carrau, Martin Beer, Sandra Blome, Alexander Schäfer

## Abstract

African swine fever virus (ASFV) poses a significant threat to pork production and wild pig populations worldwide. The study assessed the long-term fate and immunity of animals recovering from a moderately virulent ASFV infection, following the principles of a duration of immunity study for live vaccines. Pigs inoculated with the moderately virulent ASFV strain ‘*Estonia14*’ largely developed mild clinical signs and only transient viremia. Six months after the initial inoculation and once fully recovered, all the animals were challenged with highly virulent ASFV ‘*Armenia08*’. Only one of the previously exposed pigs exhibited mild clinical signs, while all control animals showed typical signs of acute, lethal ASF. Moreover, only a subset of pigs inoculated with the ASF strain ‘*Estonia14*’ displayed temporary detectability of ASFV genomes following challenge infection. Virus isolation corroborated these findings, with low levels of infectious virus in organs of previously inoculated pigs (28 days post challenge). Furthermore, monitoring of IgM and IgG kinetics enabled the analysis of humoral responses. IgG levels were sustained over the study period and increased slightly upon challenge infection. Lastly, plasma analysis revealed elevated complement factor C3a levels post inoculation and challenge in the recovered pigs, directly correlating with challenge virus presence. In contrast, both C3a and C5a levels were increased in the control group. It could be shown that complement system activation was mediated by the lectin pathway, possibly by interaction of mannose-binding lectins and ASFV particles. This study suggests that protective immunity following recovery can last at least six months. No cases of persistent or chronic disease were observed in convalescent pigs. These findings have implications for both vaccine development and assessment, as well as for disease control strategies including surveillance actions.

## 1. Introduction

African swine fever (ASF) has taken a heavy toll on wild pig populations and global pig production driven by factors such as international trade and wildlife movements. The panzootic dimensions and ASF-related mass mortalities and trade restrictions have led to widespread economic losses in associated sectors, e.g., in Europe [1], Asia [2, 3], and the Caribbean [4]. In many affected areas, the etiological agent, ASF virus (ASFV), has also caused devastating outbreaks among wild boar populations. These are often the source of spillover infections in domestic pigs raised on farms.

Current knowledge suggests that animals surviving ASFV infection can develop varying degrees of long-term immunity, ranging from partial to full protection. The induction and degree of immunity depends on factors such as strain specific virulence and immune evasion strategies (influencing the number of convalescent animals), as well as host genetics and environmental conditions. In the context of virulence and immune evasion strategies, it has been shown that infection with the naturally attenuated and moderately virulent genotype II ASFV strain ‘*Estonia14*’ causes mortality rates of 10 to 20% in domestic pigs, but up to 100% in wild boar [5], whereas infection with a highly virulent strain such as ‘*Armenia08*’ (genotype II) results in mostly 100% lethality under experimental conditions. Infection of domestic pigs with highly virulent, moderately virulent, and attenuated ASFV strains resulted in varying disease severities; ranging from fatal to transient and mild [6]. A possible reason for such observations is significant differences in immune modulation capabilities between the strains: ASFV was shown to interfere with the host immune response at various levels, such as interferon modulation, inflammation, apoptosis, antigen presentation, and cellular immunity [7]. Although ASF-recovered pigs showed no evident clinical signs of disease anymore, mild and nonspecific lesions in several organs (e.g., lung and heart) were reported [8]. However, although long-term effects such as scarred and fibrinous lesions in recovered pigs have been described, no evidence of persistent ASFV infection has been found in recent animal trials with strains of genotype I and II [5, 9].

Comprehensive knowledge about this duration of immunity against ASFV is key for disease management and control efforts, particularly in enzootic regions where recurrent outbreaks pose significant economic and animal health burdens. Additionally, understanding the kinetics of immunity onset, maintenance, and robustness is essential for vaccination strategies aimed at conferring long-lasting protection against ASFV. This is of particular importance, since continuous research efforts are made towards developing an effective oral vaccine against ASFV. Once developed and licensed for field use, large-scale vaccination campaigns would benefit from comprehensive knowledge about the duration of immunity induced by attenuated field isolates or modified live vaccine strains. Highly virulent isolates like ASFV ‘*Armenia08*’ will result in lethality rates close to 100% in both domestic pigs and wild boar and, therefore, will not elicit cellular or humoral immunity. However, detection of seropositive wild boar in Germany indicates that although pathogenesis of German ASFV isolates is similar to highly virulent ASFV ‘Armenia08’ [10], wild boar can survive and mount humoral immunity. However, the robustness of this immunity has not been assessed yet. Knowing kinetics and robustness of immunity after infection are essential to determine optimal vaccination schedules and booster frequencies to induce and maintain protective immunity within a pig population. Furthermore, data on the duration of immunity after vaccination could aid implementation of dynamic biosecurity measures. Data on immunity could be also implemented into epidemiological modeling and lead to cost-benefit analysis within risk assessment and biosecurity measures. Movement and trade restrictions might be relaxed in areas with high prevalences of immune animals, while tightened or maintained in areas with low prevalences of immune animals. In addition, relying solely on antibody detection to assess the immune status of an animal may be inadequate for biosafety as recent evidence suggests a potential disconnection between antibody levels and protection capacity. In some cases, antibody levels can wane faster than the actual capability to resist infection and *vice versa* [11].

Taking this into account, we wanted to shed light on the duration of immunity against ASFV in pigs following natural infection. We therefore assessed the long-term protective efficacy induced by a prior exposure to the moderately virulent ASFV strain ‘*Estonia14*’ against a subsequent challenge infection with the highly virulent strain ‘*Armenia08*’. Through a comprehensive analysis of immune parameters and clinical outcomes, we defined suitable assays, i.e., immunoglobulin isotype ELISAs and complement activation, to detect major immune checkpoints in the ASFV-specific immunity. Defining these immune characteristics in pigs recovered from infection with the moderately virulent strain ASFV ‘*Estonia14*’ might also lead to the definition of correlates of protection, thereby facilitating more targeted future vaccine developments. A challenge of the induced immunity after six months was chosen, because such studies are mandatory for licensing dossiers and the duration of immunity induced by vaccine strains of less than six months would not be sufficient. However, since live attenuated vaccine strains are based on common ASFV strains, definition of the duration of immunity after natural infection is essential to put vaccine strain data into a proper context.

## 2. Materials and Methods

### 2.1. Experimental design

The animal experiment consisted of two groups of 10 Large White domestic pigs each. The first group of animals arrived at the Friedrich-Loeffler-Institut (FLI), Riems, Germany, at the age of eight weeks. The second group entered the study after six months and was age-matched to the animals already in the study. Prior to transfer to the FLI, all animals were tested negative for ASFV by qPCR. All animals were obtained from the FLI breeding facility in Mariensee and given a two-week acclimatization period. The animal experiment was performed in accordance with the latest German animal welfare regulations. It was approved by the competent authority (Landesamt für Landwirtschaft, Lebensmittelsicherheit und Fischerei Mecklenburg-Vorpommern [LALLF M-V]) under the reference 7221.3-1.1-004/20 before the animals were obtained.

To enable indubitable identification of each individual, all animals received a unique ear tag: #91, #92, #93, #94, #95, #96, #97, #98, #99, #100 (first group) and #44, #46, #50, #53, #65, #66, #77, #80, #83, #84 (challenge control group). The two groups were held under similar housing conditions, but in different pens to ensure species-appropriate social behavior.

The first part of the study included oro-nasal inoculation of pigs to mimic natural infection routes without vector involvement. The first group (n = 10) was inoculated with 2 ml/pig of 2.5 × 10^4^ hemadsorbing units 50% (HAU_50_) per ml of the attenuated ASFV strain ‘*Estonia14*’ (genotype II). This virus strain had shown an attenuated phenotype in previous studies [5] and was chosen to allow survival and assessment of the duration of immunity. After six months, the surviving animals (n = 9), together with the control group (n = 10), were inoculated oro-nasally with 10 ml/pig of 1 × 10^4^ HAU50 per ml of the highly virulent ASFV strain ‘*Armenia08*’ (genotype II). The first group (animals inoculated with both viruses) will be referred to as ‘**EST + ARM**’ and the challenge control animals as ‘**ARM**’.

After inoculation, blood, serum, and saliva were collected weekly during the first month, and monthly for the following five months (days 0, 7, 14, 21, 28, 63, 92, 126, and 154 post infection). After challenge, all animals were sampled on days 0, 4, 7, 10, 14, 21, and 28. Throughout the study, animal well-being was assessed daily using a comprehensive clinical scoring (CS) system. This system allows scoring of changes in animal behavior (e.g., occurrence of appetite loss or lethargy) and appearance (occurrence of skin lesions, hematoma, or swellings) linked to disease progression ([12], modified). Animals were euthanized either upon reaching a clinical score of 10 points (moderate endpoint, [13]) or upon the development of clinical signs that were classified as intolerable. Furthermore, rectal temperatures were recorded daily and ≥ 40.0°C was defined as fever.

### 2.2. Viruses and Cells

For oro-nasal inoculation of all animals, porcine spleens from previous animal trials were used. The frozen spleen samples were homogenized in a mortar with sterile sea sand. The suspension was clarified by centrifugation and titrated on monocytes/macrophages derived from peripheral blood mononuclear cells (PBMCs) to ensure a sufficient ASFV titer to elicit infection in the animals.

For PBMC isolation, blood was obtained from healthy donor pigs that are kept at the FLI quarantine facility. Whole EDTA-blood was mixed with Hanks dextran (10% solution) at a ratio of 1:10. After an incubation of 90 min at room temperature, the PBMC-containing supernatant was collected and washed before seeding in Dulbecco′s Modified Eagle′s Medium (DMEM, supplemented with 10% fetal calf serum and 0.01% Penicillin/Streptomycin, Gibco). The fraction containing red blood cells (RBCs) was collected and diluted in 1X PBS at a ratio of 1:10 to perform hemadsorption tests (HATs). The HATs were used to a) titrate the inoculum prior to use to ensure sufficient ASFV titers for infection and b) back-titrate it afterwards as validation of the inoculum. Cells were seeded into 96-well plates at a density of 3 × 10^5^ cells/well and incubated at 37°C in presence of CO_2_ and a humidified atmosphere. After 24 h of incubation, recombinant colony-stimulating factor 2 (CSF2) was added at a concentration of 2 ng/ml to initiate differentiation of monocytes. Following a differentiation period of 24 h, the inoculum (either ‘*Estonia14*’ or ‘*Armenia08*’) was added to the cells (100 µl/well) in a 10-fold dilution series. All titrations were carried out in triplicates with four wells per dilution. After 24 h, donor-respective RBCs were added to each well at a ratio of 1:40, results were analyzed 24 h and 48 h later.

### 2.3. Sample collection and processing

To monitor kinetics of ASFV genome and ASFV-specific antibodies in pigs upon infection with moderately virulent ASFV ‘*Estonia14*’, blood and serum were taken from each animal on each sampling day. After six months, the surviving animals (group: EST + ARM) were challenged with the highly virulent ASFV ‘*Armenia08*’ along with challenge control animals (group: ARM). Blood and serum were collected on days 0, 4, 7, 10, 14, 21, and 28 after challenge.

Serum and EDTA-blood were collected from the *Vena jugularis* using aspiration collection tubes (KABE Labortechnik, Nümbrecht, Germany), while pigs were restrained with a snout sling or rope. Both matrices were directly subjected to nucleic acid extraction and subsequent qPCR. Part of the EDTA-blood was centrifuged to obtain blood plasma for subsequent analyses of soluble complement factors, i.e., C3a and C5a.

All samples were stored at -80°C for later use.

During necropsy, either scheduled for all survivors 28 days post challenge (dpc) or at the humane endpoint for all challenge control animals (7-8 dpc), the following organs were obtained: tonsil, lung, gastrohepatic lymph node (ghLN), mandibular lymph node (maLN), spleen, bone marrow (BM), blood, and serum. Around 200 mg of each organ sample was placed in a 2 ml centrifugation tube containing 1 ml of PBS and a 5 mm metal bead, then homogenized at 30 Hz for 3 min using a tissue lyzer (TissueLyzer II, Qiagen, Hilden, Germany). Tissue homogenates, as well as blood and serum were subjected to nucleic acid extraction and qPCR to assess ASFV genome loads. To investigate whether infectious virus can be found in organs of survivors and challenge controls, virus isolation was performed with homogenates of tonsil, ghLN, and spleen.

### 2.4. DNA extraction and qPCR

For extraction of nucleic acids, tissue homogenates, whole blood and serum samples were processed using the Nucleo-Mag® VET Kit (Macherey-Nagel, Düren, Germany) on a KingFisher 96 Flex platform (Thermo Fisher Scientific, Darmstadt, Germany), according to manufacturer’s instructions. For whole blood, 70 µl were used for nucleic acid extraction, while 100 µl were used of tissue homogenate or serum. A serum sample negative for ASFV-genome was included as extraction control. For qPCR, the virotype ASFV 2.0 PCR kit (Indical, Leipzig, Germany) was used. Each qPCR reaction was carried out according to manufacturer’s instructions, multiplexing ASFV genome (FAM), porcine *ACTB* as reference gene control (HEX), and a heterologous internal control to ensure inhibition-free performance (Cy5). Virus identity after challenge infection was confirmed by ‘Estonia14’-specific PCR established previously [5]. All qPCR reactions were carried out on a Bio-Rad C1000TM thermal cycler, equipped with the CFX96TM Real-Time System (Bio-Rad, Feldkirchen, Germany). ASFV genome copy numbers were calculated by utilizing a PCR standard containing a defined amount of extracted ASFV DNA (8 × 10^7^ genome copies/ml) in RSB50 buffer (50 ng/µL carrier RNA, 0.05% Tween 20, 0.05% sodium azide in RNase-free water).

### 2.5. Virus Isolation

Detection of infectious virus in tonsil, spleen, and gastrohepatic lymph nodes was carried out via HAT. However, to ensure accurate results for organ samples containing low amounts of infectious virus, a blind passage was carried out before evaluation of viral loads in organs. For blind passage, PBMCs were seeded into a 24-well plate at a density of 1.5 × 10^6^ cells/well, in presence of 2 ng/ml CSF2. After 24 h, 200 µl of each organ homogenate was added in duplicates and incubated for 72 h. The plates were frozen at -80°C for ≥ 24 h to ensure rupture of the cells, before ASFV-containing supernatant were used in HAT. Each technical replicate (n = 2) of the blind passage was subjected to HAT in technical replicates (n = 4). The results were divided into negative (—, all wells negative), weakly-positive (•, up to 4 wells positive), positive (••, 4-8 wells positive), and strongly positive (•••, 4-8 wells positive, high rousette counts).

### 2.6. Serology

To monitor kinetics of ASFV-specific antibodies, all serum samples were subjected to several serological assays. First, all samples were evaluated using two routinely used and accredited ELISA kits: (I) ID Screen® ASF Indirect (ID.vet, Grabels, France), detecting antibodies targeting ASFV p32, p62, p72, and (II) Ingezim PPA COMPAC (Gold Standard Diagnostics, Freiburg, Germany), detecting p72 targeting antibodies. All assays were performed in accordance with manufacturer’s instructions. However, since these assays either only detect porcine IgG or do not distinguish between Ig isotypes at all, an indirect ELISA protocol was additionally employed to not only evaluate whole ASFV-specific immunoglobulins, but to assess IgG and IgM kinetics individually. Therefore, ASFV-antigen was obtained from the European Union Reference Laboratory for ASF (EURL-ASF), CISA-INIA (Spain). Microtiter plates (medium-binding plate, Greiner) were coated with ASFV-antigen in sodium carbonate buffer, as described in the SOP provided by the EURL-ASF (SOP/CISA/ASF/ELISA/1, REV.5, 2021). In brief, all plates were washed after incubation overnight, dried and stored at -20°C until further use. To reduce unspecific binding, all plates were incubated with a blocking solution (1X PBS supplemented with 5% horse serum) for 3 h at RT. All sera were diluted with washing solution (1X PBS supplemented with 0.05% Tween-20 (Roth, Karlsruhe, Germany)) at a ratio of 1:30 immediately prior to performing the experiment. Reference sera from the German National Reference Laboratory for ASF (NRL-ASF) were included on each plate as positive and negative controls. For normalization purposes, two wells on each plate were not coated with ASFV-antigen and only treated with sodium carbonite coating buffer. After incubation and subsequent washing, the respective, HRP-conjugated secondary antibody was added to each well: for (I) IgG detection, a polyclonal anti-porcine IgG antibody (#A100-205P, Biomol, Hamburg, Germany) was added at a dilution of 1:10,000, whereas for (II) IgM detection, a polyclonal anti-porcine IgM antibody (#AAI48, Bio-Rad, Feldkirchen, Germany) was added at a dilution of 1:5,000. The signal was acquired with an ELISA reader (TECAN, Switzerland).

Activated complement factors C3a and C5a were measured using species-specific and commercially available kits according to manufacturer’s instructions (MyBioSource, San Diego, USA). Evaluation of C3a and C5a levels was conducted with plasma of the pigs, as plasma samples are more accurate in detecting soluble biomarkers compared to serum samples [14]. For *in vitro* assays, untreated serum from animals of this study taken 0, 7, and 154 dpi was mixed 1:2 with either PBS, 10 mM EDTA, or 1 mM EGTA/2 mM MgCl_2_ (MgEGTA) and incubated for 30 min at 37 °C. Afterwards, the treated serum samples were mixed with 10^6^ HAD_50_/ml ‘*Armenia08*’ and incubated for 90 min at 37 °C. The reaction was stopped by addition of 10 mM EDTA and 50 µg/ml Futhan (BD Biosciences, Heidelberg, Germany). Levels of C3a and C5a were investigated by C3a and C5a ELISA as described above. For analysis of complement-dependent infection rates, untreated or heat-inactivated (56 °C, 30 min) plasma samples of six naïve animals were incubated with 10^6^ HAD_50_/ml particles of either ASFV ‘*Estonia14*’ or ASFV ‘*Armenia08*’ for 90 min at 37 °C. The reaction was stopped by addition of 10 mM EDTA and 50 µg/ml Futhan (BD Biosciences, Heidelberg, Germany). Pretreated ASFV particles were used for infection of porcine monocytes (prepared 48 h in advance). Infection rates were assessed 48 h later by indirect immunofluorescence with anti-ASFV-p72-FITC (polyclonal rabbit serum, in-house) and DAPI counterstain on a Leica Thunder Imager (Leica Microsystems, Wetzlar, Germany). Using the automated event-counting function of ImageJ, all cells/well were counted (all DAPI-positive events and the ration of infection was calculated by counting and normalizing infected cells (FITC-positive events) to total cell counts.

### 2.7. Statistics

Statistical analyses and data visualization were performed with GraphPad Prism 10.3.1 for Windows (GraphPad Software Inc., Boston, USA). Sample numbers are indicated in the figure legends. Differences in survival and clinical scores between previously *‘Estonia14’*-infected and control animals were assessed by unpaired *t*-tests of the area under the curve (AUC). Significant differences in complement activation were analyzed by a 2-way-ANOVA with Holm-Šidák’s correction for multiple comparisons. Significant statistical differences are indicated by their *p*-value.

## 3. Results

### 3.1. Course of ASFV ‘Armenia08’ infection in pigs previously infected with ASFV ‘Estonia14’

After inoculation with ASFV ‘*Estonia14*’, all 10 animals developed fever and first clinical signs of disease 4 to 5 dpi (Figure 1). The body temperature was elevated in all animals 5 dpi ranging from 39.5°C to 41.8°C until 10 dpi (Figure 1A). Clinical signs of disease were mild in all animals in the first week of infection, where lethargy, loss of appetite, and reduced liveliness could be observed (Figure 1B). All but one animal fully recovered until 14 dpi, with no elevated body temperature or clinical signs. One individual (#94) succumbed to infection and was found dead in the pen at 9 dpi. The ASFV genome loads in ‘*Estonia14*’-infected animals were comparable in all animals in the first 4 weeks after inoculation (Figure 1C). The detected ASFV genome loads were highest 7 dpi with 18.4±0.9 and decreased slowly to 21.7±1.0 28 dpi. During the next months, the animals were sampled once a month. A steady decrease of ASFV genome loads was observed, with Cq values of 27.3±1.2 after two months and 32.5±1.7 after three months. The first animal (#95) was ASFV-negative at this time. By month 4, only animal #92 was still ASFV-positive in blood and all animals were negative five months after infection.

**Figure 1.**
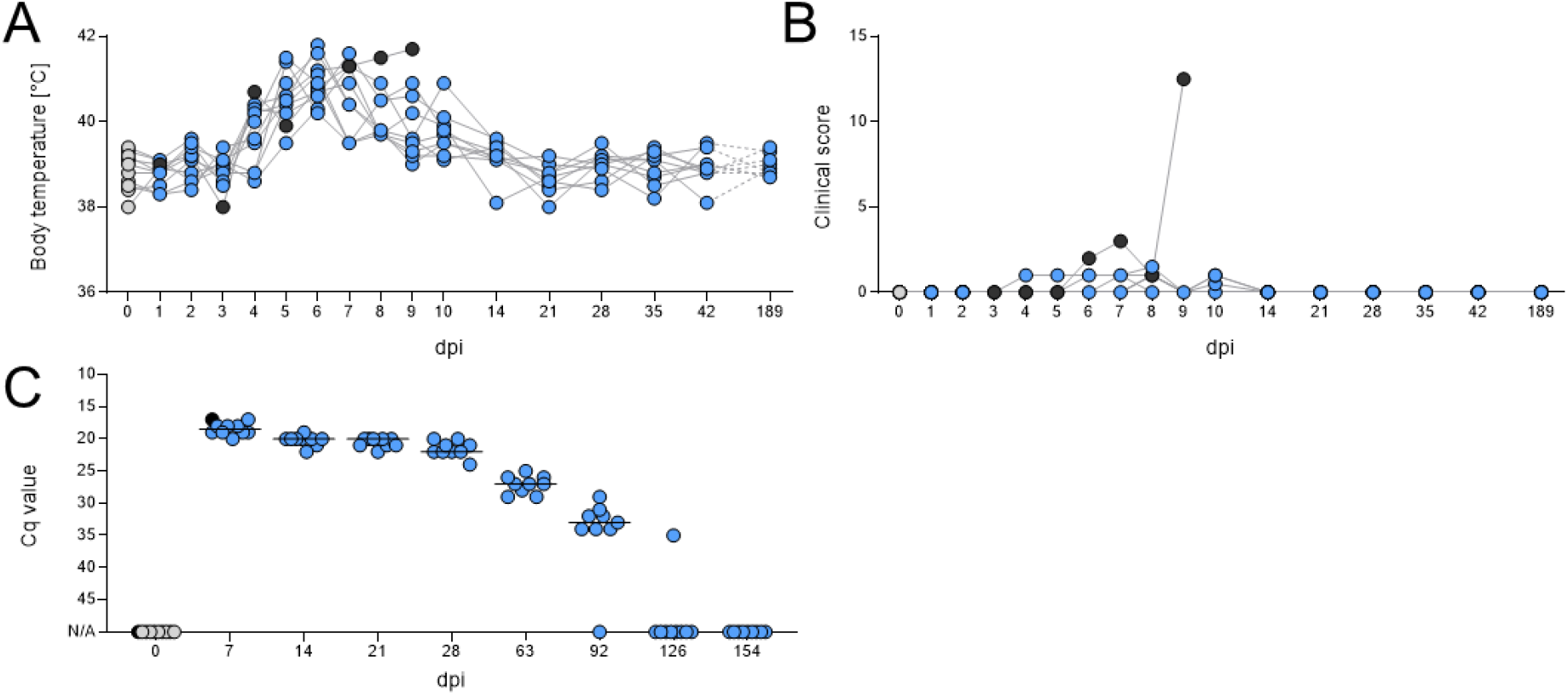
(**A**) Body temperature, (**B**) clinical score, and (**C**) ASFV genome loads in blood of pigs (*n* = 10) upon inoculation with ASFV ‘*Estonia14*’. Black dots show one individual (#94) that developed severe clinical symptoms and succumbed to infection 9 dpi. Lines represent medians, dots individual animals.

After an observation period of six months, all remaining pigs, along with a control group, were inoculated with ASFV ‘*Armenia08*’ to assess the robustness of the ASFV-specific immunity. Only one individual (#93) in the EST+ARM group developed fever after challenge infection, however, the fever did not persist (Figure 2A). The clinical score of this animal on the days of fever included scores for gait and bearing, possibly resulting from bacterial infection between the claws, not an ASFV-specific manifestation. In contrast, all animals of the ARM group developed high fever, ranging from 40°C 5 dpi to 41.8°C 8 dpi (Figure 2B). Onset of fever in the control animals was accompanied by a disease manifestation typical for ASFV infection: severe lethargy, no appetite, dehydration, pain in extremities with difficulties standing/walking, and respiratory distress. All control animals reached the humane endpoint 7-8 dpi. Cq-values in the ARM group ranged from 19.2±1.7 4 dpc to 15.3±0.4 on the day of euthanasia (7/8 dpc). Among the EST+ARM group, only three animals (#92, #95, #98) remained ASFV genome-negative in blood throughout the observation period after challenge infection with ASFV ‘*Armenia08*’, while the remaining six animals were ASFV-positive at least once (Figure 2C). However, Cq values were considerably lower than in control animals and ranged from 37.2±0.5 7 dpc to 31.1±5.1 on the last day, 28 dpc.

**Figure 2.**
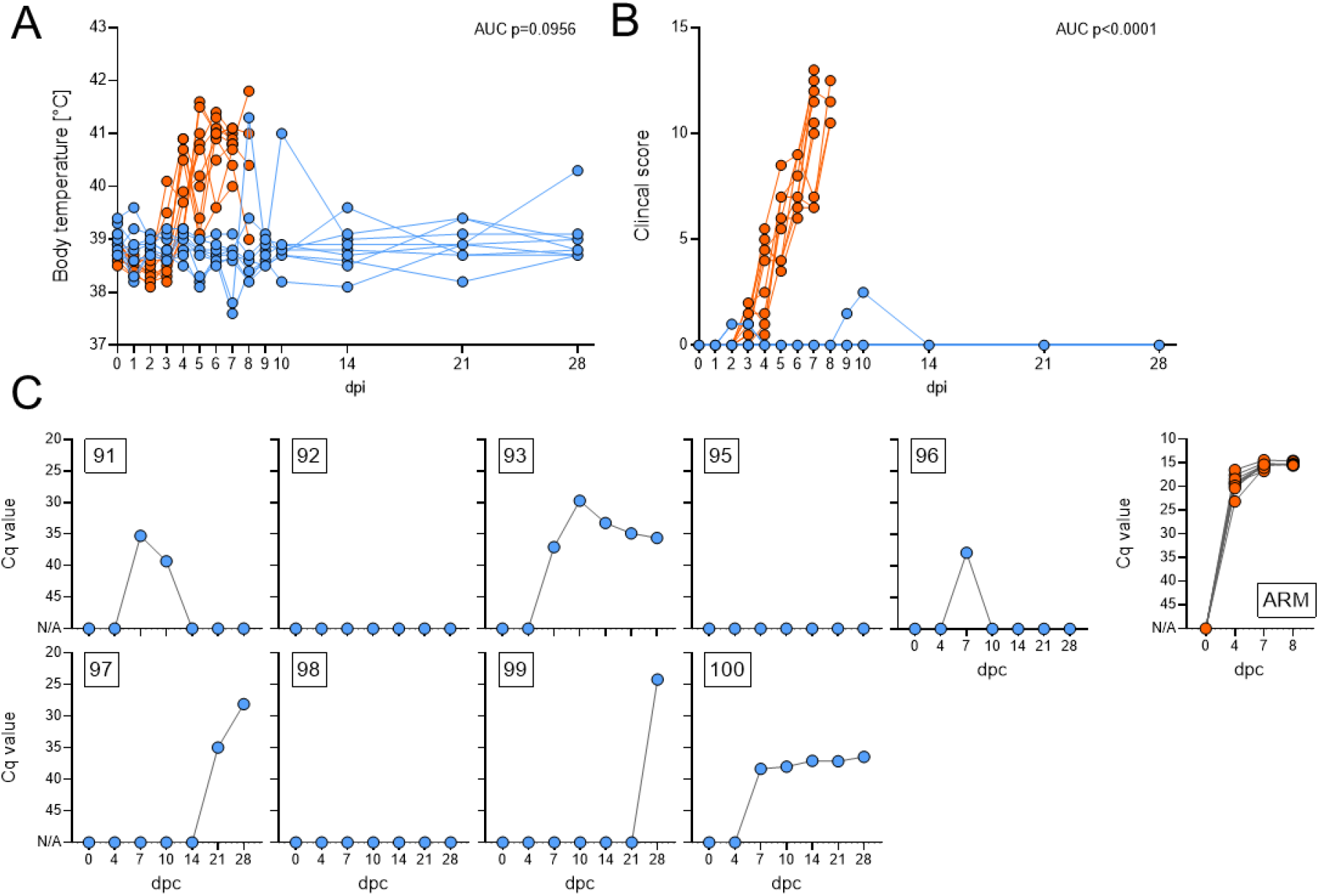
Body temperature and clinical score of (**A**) previously ‘*Estonia14*’-inoculated (*n* = 9, blue) and (**B**) naïve pigs (*n* = 10, orange) after challenge with ASFV ‘*Armenia08*’. (**C**) ASFV genome loads in blood of challenged animals. Positive results in the blood were verified as ASFV ‘Armenia08’ challenge virus by ASFV ‘*Estonia14*’-specific qPCR. Numbers indicate ear tags.

Gross pathological findings in the ARM group and #94 of the EST+ARM group comprised of typical manifestations during ASFV infection: hemorrhages in several organs (e.g., petechiae in kidneys), ascites, and pericardial effusion. In contrast, gross pathology in the EST+ARM group rendered no abnormalities, except scarred and fibrinous lesions in and around the pericardium of #93, #97, #99, and #100, indicating a previous pericardial effusion or bacterial colonialization. The ASFV genome loads were assessed in several organs (Figure 3). In the ARM group, mean ASFV genome copy numbers per run were consistent between animals, being 4.16E+04 in tonsil, 3.03E+04 in lung, 4.13E+04 in gastrohepatic LN (ghLN), 3.58E+04 in mandibular LN (maLN), 2.27E+05 in spleen, 9.93E+03 in bone marrow (BM), and 1.44E+04 in serum. In contrast, only #93, #97, #99, and #100 of the EST+ARM group were weakly positive in organs at the time of euthanasia. The ASFV genome copy numbers in the EST+ARM group were 2.6E+00 in tonsil, 1.13E+00 in lung, 8.34E+01 ghLN, 4.22E+01 in spleen, and 1.24E+01 in serum. No ASFV genomes were detected in maLN and BM. Of note, ASFV genome copies in those animals originated from the challenge virus, as an ASFV ‘*Estonia14*’-specific qPCR was negative for blood and organ samples.

**Figure 3.**
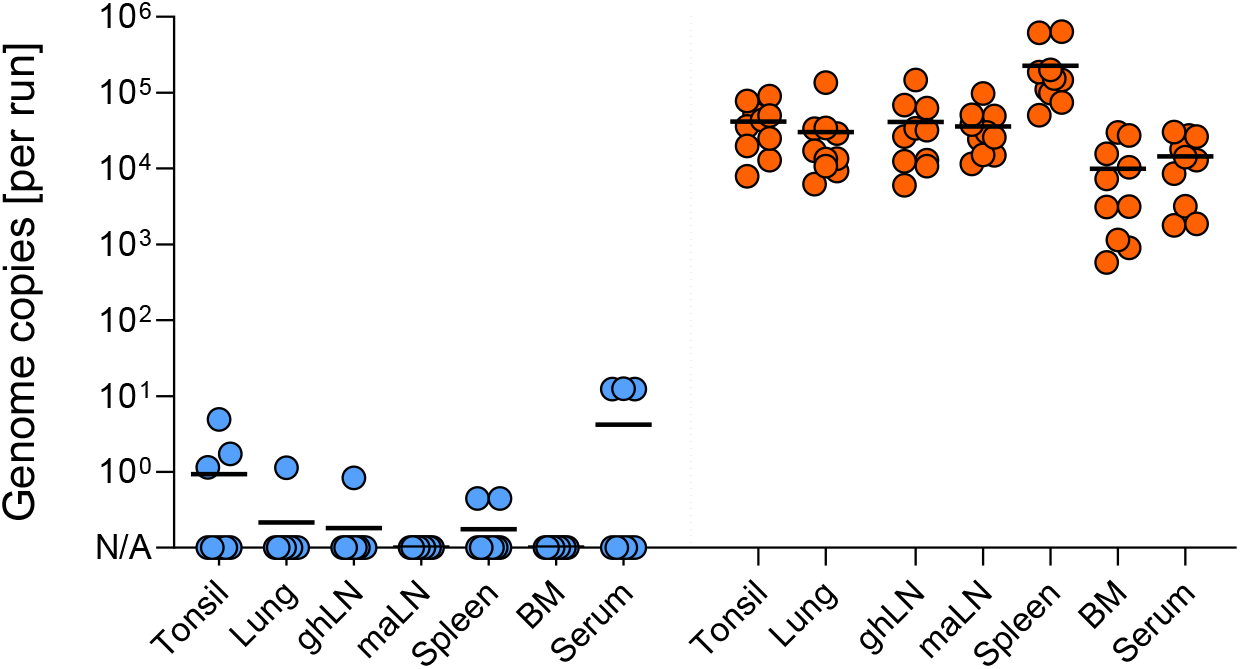
Detection of ASFV genome after challenge with ASFV ‘*Armenia08*’ in organs of pigs previously infected with ‘*Estonia14*’ (*n* = 9, blue dots, left, 28 dpc) or naïve animals (*n* = 10, orange dots, right, 7-8 dpi). Lines represent medians, dots individual values. ghLN, gastrohepatic lymph node; maLN, mandibular lymph node; BM, bone marrow.

Subsequently, tonsil, gastrohepatic LN, and spleen samples were subjected to virus isolation (Table 1). High amounts of infectious virus particles were found in all animals of the ARM group, as well as in animal #94 of the EST+ARM group. Moderate amounts of infectious particles were found in tonsil and ghLN samples of #93, #97, #99, and #100, while spleen samples were weakly positive.

**Table 1.** Virus isolation results in organs of pigs previously infected with ‘*Estonia14*’ (EST+ARM) or naïve animals (ARM)

| EST+ARM |  |  |  | ARM |  |  |  |
| --- | --- | --- | --- | --- | --- | --- | --- |
|  | Tonsil | ghLN | Spleen |  | Tonsil | ghLN | Spleen |
| #91 | — | — | — | #44 | •• | ••• | ••• |
| #92 | — | — | — | #46 | •• | ••• | ••• |
| #93 | •• | •• | •• | #50 | •• | ••• | ••• |
| **#94** | ••• | ••• | ••• | #54 | ••• | ••• | ••• |
| #95 | — | — | — | #65 | ••• | ••• | •• |
| #96 | — | — | — | #66 | ••• | ••• | ••• |
| #97 | •• | •• | • | #77 | ••• | ••• | •• |
| #98 | — | — | — | #80 | ••• | ••• | ••• |
| #99 | •• | • | • | #83 | •• | ••• | ••• |
| #100 | •• | •• | • | #84 | ••• | ••• | •• |

— = negative (0/8); • = weakly positive (1-4/8); •• = positive (5-8/8); ••• = highly positive (8/8 with high rousette count), ghLN, gastrohepatic lymph node
**Bold** = animal succumbed to ASFV ‘*Estonia14*’ infection 9 dpi

### 3.2. Kinetics of IgM and IgG in serum of pigs recovered from ASFV infection

To assess the humoral response in EST+ARM pigs, ASFV-specific IgM and IgG kinetics were monitored (Figure 4). Levels of IgM were highest 7 and 14 days after inoculation with ‘*Estonia14*’ and, although declining, remained above threshold in most animals. Upon challenge with ‘*Armenia08*’, levels of IgM increased again. However, levels were lower compared to day 7 and 14 of ‘*Estonia14*’ infection and declined rapidly. First detection of ASFV-specific IgG in all animals was possible 14 days after inoculation with ‘*Estonia14*’. IgG titers increased over time and remained relatively stable between months 2 and 6 after inoculation. Upon challenge, IgG levels moderately increased 14 and 21 dpc, but were comparable to 0 dpc at the end of trial (28 dpc). These results, generated with an ELISA against whole virus lysate, rendered comparable results to a commercially available competitive ELISAs, targeting the late protein p72 or the early protein p30/p32 (Supplementary Figure 1). However, the commercially available ELISAs could not differentiate between Ig isotypes and failed to detect changes in IgM and IgG levels after challenge infection.

**Figure 4.**
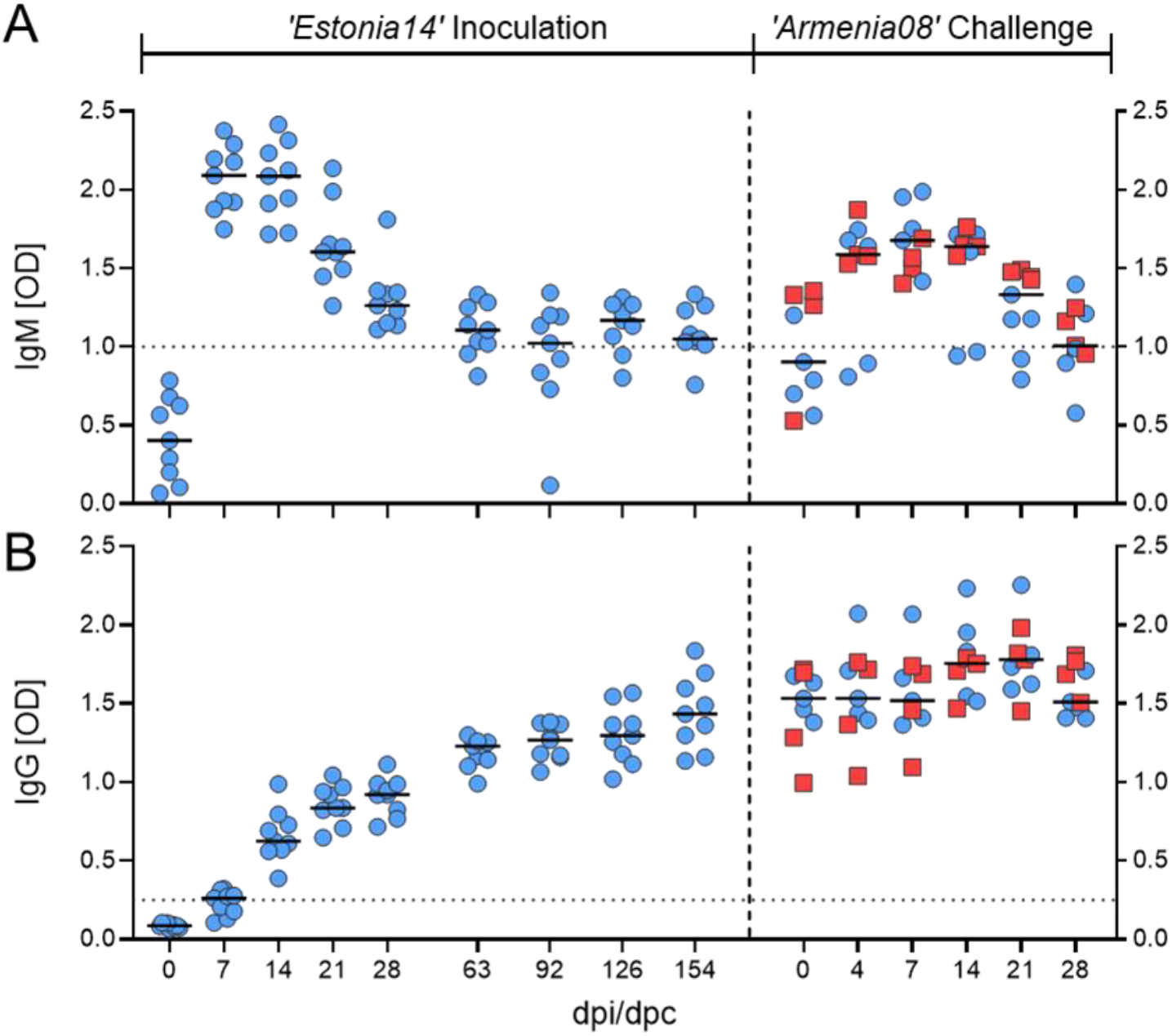
Kinetics of (**A**) IgM and (**B**) IgG in sera of pigs after inoculation with ASFV ‘*Estonia14*’ and challenge with ASFV ‘*Armenia08*’ (*n* = 9). Animals that were still positive in qPCR at the end of the trial (28 days after challenge) are shown as red squares. Lines represent medians, dots individual values.

### 3.3. Complement activation upon infection with moderately and highly virulent ASFV in recovered and naïve pigs

To further evaluate soluble inflammation factors, plasma was collected on all sampling days and subjected to species-specific investigation of complement activation by detection of complement components 3 (C3a) and 5 (C5a) in all animals (Figure 5A, B). During ‘*Estonia08*’ infection, C3a levels significantly increased in EST+ARM animals 14 dpi. After ‘*Armenia14*’ challenge, C3a levels in EST+ARM animals increased slightly but only reached significance at 28 dpc (Figure 5A). C5a levels remained unaffected during both infections in the EST+ARM group. In ARM animals, levels of C3a significantly increased during ‘*Armenia14*’ infection, as early as 4 dpc. Contrary to EST+ARM animals, the levels of C5a were significantly increased in ARM animals 7 dpc (Figure 5B).

**Figure 5.**
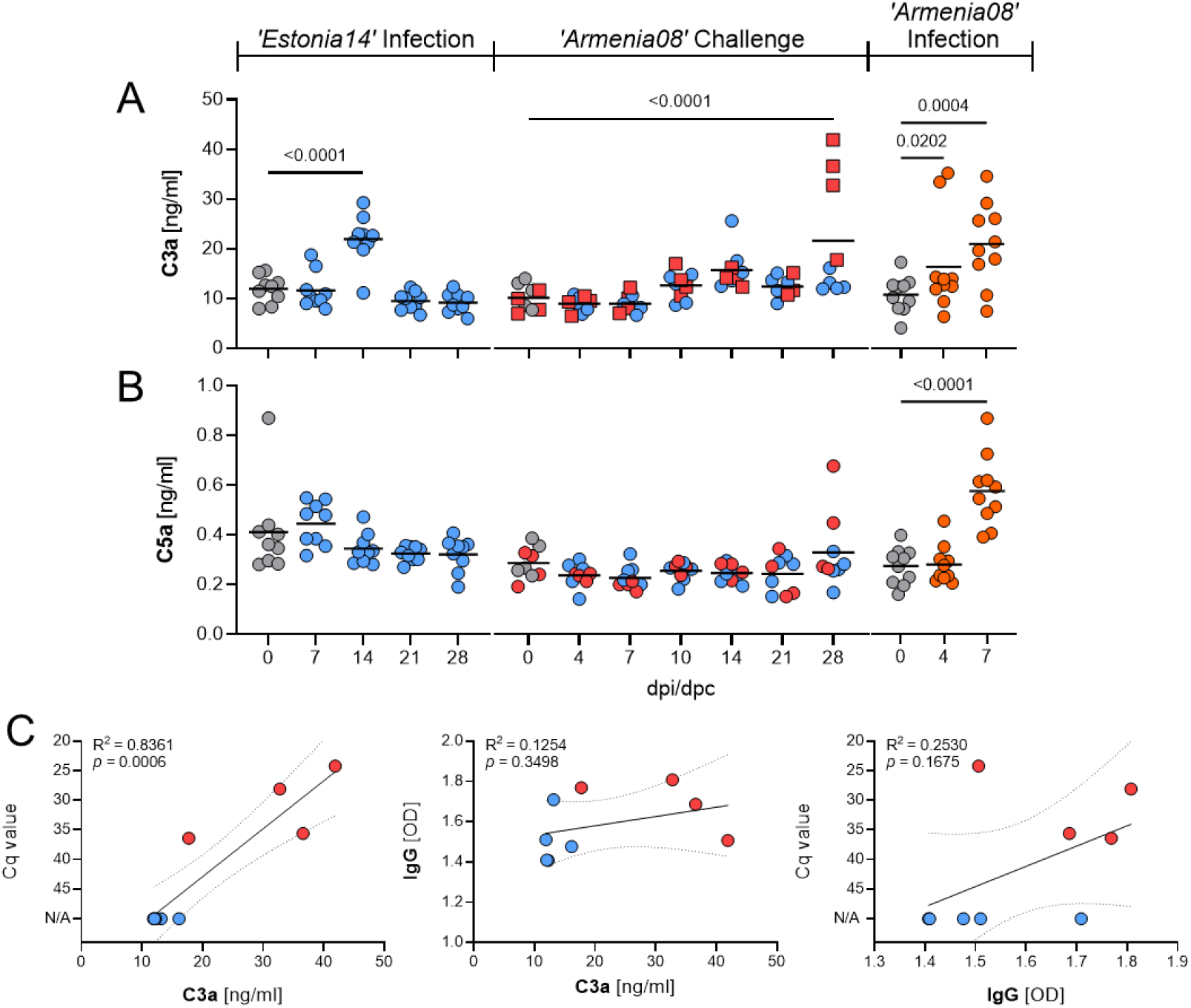
Detection of activated complement components (**A**) C3a and (**B**) C5a in plasma after infection with ASFV ‘*Estonia14*’, challenge with ASFV ‘*Armenia08*’ (*n* = 9, blue dots), or after infection in control animals (orange dots, *n* = 10). The respective reference days (day 0 before infection/challenge) are displayed in grey. Lines represent medians, dots show individual values. Red squares among EST+ARM animals indicate individuals where live virus was isolated 28 dpi. Significance was calculated by 2-way-ANOVA with Holm-Šidák’s correction for multiple comparisons. **(C)** Correlation analysis in EST+ARM animals.

Of note, the four animals with the highest C3a levels among EST+ARM animals after challenge infection were animals which still contained infectious virus in organs at the end of the trial (#93, #97, #99, #100). Cq-values in EST+ARM animals were positively correlated with C3a levels 28 dpc (R^2^=0.8361, *p*=0.0006; Figure 5C). In contrast, IgG titers were not predictive of C3a levels and Cq values did not correlate with IgG levels.

Additionally, we investigated the mechanism behind complement activation in additional *in vitro* assays. Serum of pigs at different stages after ‘*Estonia14’* infection was incubated directly with ASFV ‘*Armenia08’*. The stages were chosen based on the presence of serum antibodies to assess the classical complement pathway: naïve serum (0 dpi) contained no antibodies, early serum (7 dpi) contained IgM but little to no IgG, and late serum (154 dpi) contained high levels of IgG (Figure 4A, B). In order to identify the complement pathway that led to activation, we further added either PBS (control, no inhibition), EDTA (complete complement inhibition), or EGTA/MgCl_2_ (MgEGTA, specific inhibition of classical and lectin pathway) to the serum. We found a significant decrease of C3a and C5a levels after incubation with EDTA and MgEGTA (Figure 6A, B). Subsequently, we asked whether decoration of ASFV particles by complement factors would increase viral uptake and infection rates. Viral particles of ‘*Estonia14*’ or ‘*Armenia08*’ were incubated with untreated or heat-inactivated plasma from naïve animals and used for *in vitro* infection of porcine monocytes for 48 h. Infections rates were assessed by indirect immunofluorescence. We found a significant increase when particles were incubated with heat-inactivated plasma, irrespective of viral strain used (Figure 6C).

**Figure 6.**
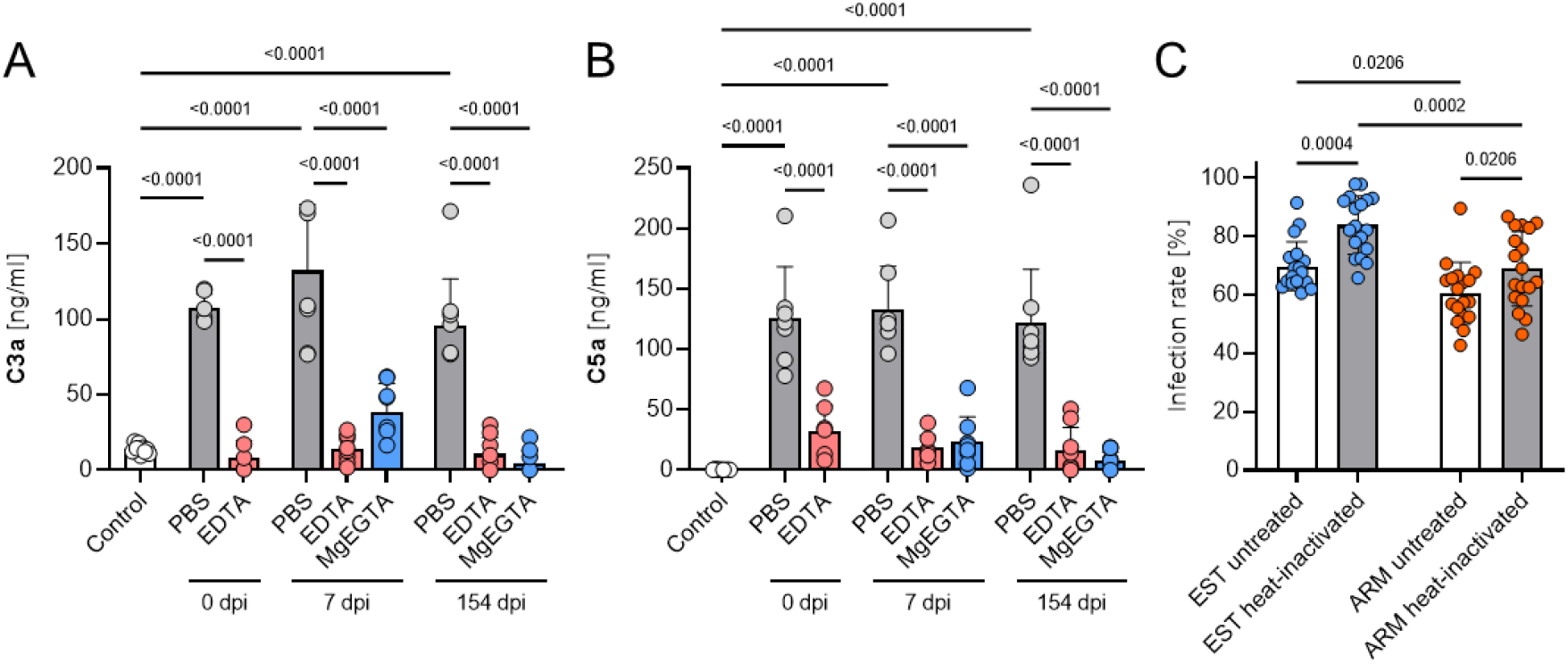
Detection of activated complement components (**A**) C3a and (**B**) C5a *in vitro*. Untreated serum of the indicated times after ‘*Estonia14’* infection (plasma of 7 and 154 dpi contains ASFV-specific IgM/IgG) was mixed with PBS (grey), EDTA (red), or EGTA/MgCl_2_ (MgEGTA, blue) and then incubated with 10^6^ HAD_50_/ml ASFV ‘*Armenia08’*. (**C**) Infection rates of monocytes 48 h after infection with ‘*Estonia14*’ (blue) or ‘*Armenia08*’ (red) particles that were pre-incubated with either untreated (white bars) or heat-inactivated (grey bars) plasma from naïve animals (*n* = 6) for 90 min at 37 °C. Significance was calculated by One-way-ANOVA with Holm-Šidák’s correction for multiple comparisons.

## 4. Discussion

Long-term studies on potential protective immunity against ASFV infections provide important data for duration of immunity, improved vaccine designs and potential management option for this notifiable disease. However, these studies are challenging, due to their high demands to animal husbandry within high-containment research facilities. In the present study, inoculation with the moderately virulent ASFV ‘*Estonia14*’ induced protective immunity against the highly virulent ASFV ‘*Armenia08*’ for at least six months in domestic pigs. Interestingly, although all animals were protected from clinically apparent disease, viral genomes were still detected and live virus was isolated from some animals after challenge infection, although titers of recovered ASFV were not assessed.

Previous studies with various ASFV strains demonstrated varying degrees of protection against lethal challenge infection. Inoculation with the naturally attenuated ASFV *‘OURT88/3’* did not convey protective immunity against highly virulent *‘Benin 97/1’* at 130 days after inoculation [15], contrasting the robust and protective immunity for at least six months (189 days) observed in our study. These observations possibly indicate the development of strain-specific immunity, varying in duration and strength. Such differences may result from strain-specific variations in the immunogenicity of ASFV antigens and capsid-specific antibody responses elicited by different ASFV strains, ultimately altering humoral immune responses [16, 17]. It has been demonstrated that such differences in antibody responses occur naturally between breeds of domestic pigs and even between individuals within a breed [17]. These conclusions were further validated by observing no such differences in the capsid-specific antibody response in Babraham pigs after inoculation with the moderately virulent ASFV strain ‘*OURT88/3*’ [18]. Babraham pigs are inbred and express homozygous SLA molecules, excluding the influence of individual genetic and immunological factors [19]. However, although some novel antigens were introduced with the ‘*Armenia08*’ strain in the 5’-region of the viral genome with the challenge infection ([5], evidenced by increasing IgM levels in EST+ARM pigs after challenge), strain-specific immunity conveyed by ASFV ‘*Estonia14*’ was sufficient to protect from a different highly virulent strain, ASFV ‘*Armenia08*’.

Furthermore, host genetics and the environmental status of pigs can significantly impact disease outcome. For example, inoculation of domestic pigs and wild boar with ASFV ‘*Estonia14*’ under standardized conditions resulted in no lethality in domestic pigs and 60% lethality in wild boar [5]. These differences in disease manifestation and outcome may be due to distinct response patterns of CD8+ cytotoxic T cells in domestic pigs and wild boar [20]. The environmental conditions in housing facilities of pigs also influences the disease outcome: upon inoculation with ASFV ‘*Estonia14*’, SPF (specific pathogen free) pigs had reduced capacity to control early virus replication, but presented overall milder disease with full protection and recovery compared to conventionally housed pigs [21]. Although recovered pigs showed no evident clinical signs of disease anymore, mild and nonspecific lesions in several organs (e.g., lung and heart) were reported [8]. Although long-term effects such as scarred and fibrinous lesions in recovered pigs have been described, no evidence of persistent ASFV infection has been found to date [9]. However, the data of our trial indicates that immune pigs do not develop clinical signs upon challenge infection. In addition to the finding of ASFV genomes in the blood and the spleen of some immune animals, infectious ASFV particles were also found (no quantification of titers was carried out). Whether the amount of challenge virus shed by clinically inapparent pigs would suffice to infect naïve penmates remains to be determined. The possible impact of such asymptomatic carriers in wild boar populations might be detrimental and needs to be addressed in future risk assessment discussions.

A more comprehensive characterization of antiviral responses in ASFV-infected pigs is pivotal for an improved understanding of ASF pathogenicity and host responses. Detection of disease correlates in easily accessible matrices like serum could increase our understanding of ongoing infections. We found increased levels of complement factors C3a but not C5a in ASFV ‘*Estonia14*’-infected animals. Challenge infection with ‘*Armenia08*’ of recovered animals resulted in similar responses, but only in animals that were found to shed live challenge virus. In contrast, infection with ‘*Armenia08*’ in naïve pigs resulted in a concurrent increase of both C3a and C5a levels.

The complement system can be activated by three distinct pathways: the classical pathway, using antibody complexes on the target membrane, the lectin pathway, using mannose-binding lectin (MBL) detecting specific membrane-bound carbohydrates, and the alternative pathway, which is activated by spontaneous hydrolysis of complement factor C3. The complement system also reacts to viral infections for which many viruses evolved escape strategies [22]. To further characterize the involvement of complement during ASFV infection, we conducted *in vitro* experiments by specifically inhibiting or excluding all or some complement pathways: EDTA, inhibiting all complement pathways, MgEGTA, specifically inhibiting the classical and lectin pathways [23], and serum containing no ASFV-specific antibodies, excluding the classical pathway. We found a significant increase of both C3a and C5a when serum of naïve and recovered animals was incubated with ASFV. This was blocked by MgEGTA, excluding the alternative pathway. Samples taken 0 dpi, containing no ASFV-specific antibodies, also showed no complement induction, indicating that the classical pathway is not the main contributor. This leaves the lectin pathway as the inducer of complement responses during ASFV infections. While MBL is often thought to specifically bind to bacterial carbohydrates, it has been shown to detect a variety of viruses, including human immunodeficiency virus (HIV [24]), hepatitis B virus (HBV [25]), hepatitis C virus (HCV [26]), SARS-CoV-2 [27], human cytomegalovirus/ human herpesvirus-5 (CMV/HHV-5 [28]), Ebola virus (EBOV [29]), adenovirus 2 and 5 [30], Chandipura virus (CHPV [31]), Dengue virus [32], and West Nile virus (WNV [33]) in part by binding directly to viral surface proteins. The effects of complement activation during infection might be anti-as well as proviral [22, 34]. Increased C3a levels are also indicative of heightened levels of C3b, responsible for opsonization of pathogens and might thus contribute to increased uptake of opsonized particles in myeloid cells. However, heat-inactivation of the complement system led to increased infection rates of macrophages, indicating that at least some viral particles are neutralized by the activated complement system.

Independent of the immunological processes behind the observed variations, understanding the robustness and duration of immunity after infection with ASFV is essential for advancing disease management, control strategies, and risk assessment in both enzootic and non-enzootic regions. The disease has a major economic impact on the pork industry due to high fatality rates, indiscriminate culling, and trade restrictions installed to prevent viral spread. If immunity after ASFV infection or vaccination proves to be robust and long-lasting, it could fundamentally reshape control measures. For instance, evidence of a durable immunity could support more targeted depopulation policies. A satisfactory number of successfully vaccinated animals could lead to a reduction of the need for indiscriminate culling, ultimately minimizing economic losses. On the other hand, if immunity even after vaccination is incomplete or short-lived, stringent biosecurity measures like rapid diagnostics, and whole farm cullings remain a necessity. Furthermore, a reduced duration of immunity might also influence future vaccination concepts like booster intervals.

In Europe, studies for identifying the duration of immunity after vaccination are mandatory to be included in a dossier for licensing a vaccine candidate. Once the data is collected and a vaccine is fully licensed, knowing how long the induced immunity will convey protection is crucial for the development of vaccination strategies. Given that the duration of immunity is at least six months, but varies between strains (and possibly ASFV vaccine strains as well), large-scale vaccination campaigns for wild pigs can be optimized and refined, especially the regimen for booster vaccinations or the need for multi-component vaccines targeting different viral mechanisms. It is important to understand whether natural infection provides lasting immunity in order to guide the design of vaccines, which mimic such protection. The data about the duration of immunity after inoculation with an attenuated ASFV field strain or a vaccine strain can be implemented into epidemiological modeling algorithms to predict how many boosters at what time frames will be necessary to sufficiently immunize a wild population and hamper viral spread. Such modeling could predict how and if ASFV sustains itself after populations were immunized. Additionally, if animals lose immunity over time, populations may become susceptible to reinfection, necessitating the reevaluation of biosecurity timelines and quarantine durations, particularly in high-risk zones.

## 5. Conclusions

Our study investigated the duration and robustness of immunity induced by a moderately virulent ASFV strain, ‘*Estonia14*’, against a related highly virulent strain, ‘*Armenia08*’. Pigs that survived an infection with ‘*Estonia14*’ exhibited no clinical signs upon challenge infection with ‘*Armenia08*’, while control animals developed acute disease. However, some ‘*Estonia14*’-inoculated pigs showed limited viral presence post-challenge. Stable IgG levels post-inoculation and a moderate increase post-challenge further support the role of the specific humoral immunity. Interestingly, elevated complement factor C3a levels correlated with challenge virus presence in ‘*Estonia14*’-inoculated pigs, suggesting involvement of the complement system in ASFV manifestation. Conversely, increased C3a and C5a levels in control animals may indicate contribution of the complement system to ASF pathogenesis. These findings demonstrate that prior infection with a moderately virulent ASFV strain elicits robust protection against infection with a highly virulent strain for at least six months.

## Supplementary Materials

Figure S1: Two commercially available ELISA were used as comparison for the full virus antigen ELISA.

## Author Contributions

Conceptualization, S.B., M.B., A.S. and V.F.; methodology, V.F. A.S., S.B., T.C., K.W. and P.D.; formal analysis, V.F., A.S., S.B.; investigation, V.F., T.C., P.D.; data curation, V.F., A.S.; writing—original draft preparation, V.F., A.S., S.B.; writing—review and editing, V.F., A.S., S.B., K.W. and M.B.; visualization, V.F., A.S.; supervision, A.S., S.B., M.B.; project administration, S.B.; funding acquisition, S.B. All authors have read and agreed to the published version of the manuscript.

## Institutional Review Board Statement

This research was conducted in accordance with German animal welfare regulations, including EU Directive 2010/63/EC and institutional guidelines. Approval was received by the State Office for Agriculture, Food Safety and Fishery in Mecklenburg—Western Pomerania (LALFF M—V) and is filed under reference number 7221.3-1.1-004/20.

## Informed Consent Statement

Not applicable.

## Data Availability Statement

All data is available upon request from Sandra Blome, corresponding author.

## Acknowledgments

We would like to thank all animal caretakers: Matthias Jahn, Domenique Lux, Patrice Mary and Steffen Brenz for excellent animal husbandry and assistance during sampling procedures. Additionally, we would like to thank Christian Loth and Ralf Redmer for preparing and assisting all necropsies. For excellent technical assistance during sample processing and subsequent experiments, we thank Ulrike Kleinert and Stefanie Knöfel.

## Supplementary Figure 1

**Figure S1.**
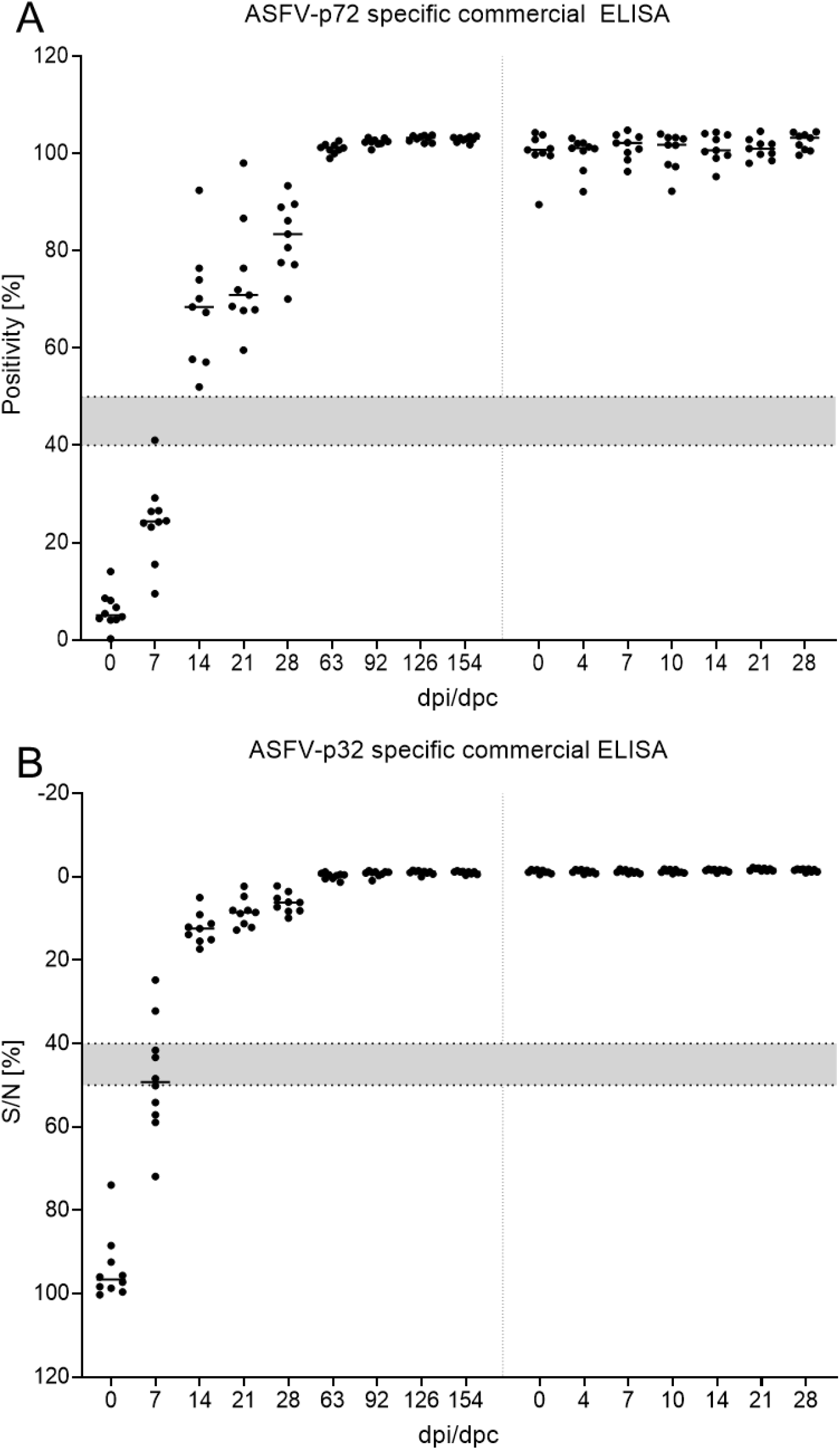
Two commercially available ELISA kits were employed to detect (**A**) ASFV-p72- and (**B**) ASFV-p32-specific antibodies in serum of EST+ARM pigs. Points represent individuals, lines represent the median. Grey areas within each plot mark the range of questionable results. All points within grey areas were interpreted as ‘questionable’, all points below as ‘negative’, all points above as ‘positive’.

## References

1. European Food Safety, A., et al., Epidemiological analysis of African swine fever in the European Union during 2022. EFSA J, 2023. 21(5): p. e08016.

2. Nguyen-Thi, T., et al., An Assessment of the Economic Impacts of the 2019 African Swine Fever Outbreaks in Vietnam. Front Vet Sci, 2021. 8: p. 686038.

3. You, S., et al., African swine fever outbreaks in China led to gross domestic product and economic losses. Nat Food, 2021. 2(10): p. 802–808.

4. Jean-Pierre, R.P., A.D. Hagerman, and K.M. Rich, An analysis of African Swine Fever consequences on rural economies and smallholder swine producers in Haiti. Front Vet Sci, 2022. 9: p. 960344.

5. Zani, L., et al., Deletion at the 5’-end of Estonian ASFV strains associated with an attenuated phenotype. Sci Rep, 2018. 8(1): p. 6510.

6. Gallardo, C., et al., Dynamics of African swine fever virus (ASFV) infection in domestic pigs infected with virulent, moderate virulent and attenuated genotype II ASFV European isolates. Transbound Emerg Dis, 2021. 68(5): p. 2826–2841.

7. Wu, L., et al., Regulation and Evasion of Host Immune Response by African Swine Fever Virus. Front Microbiol, 2021. 12: p. 698001.

8. Lai, D.C., et al., The study of antigen carrying and lesions observed in pigs that survived post African swine fever virus infection. Trop Anim Health Prod, 2022. 54(5): p. 264.

9. Petrov, A., et al., No evidence for long-term carrier status of pigs after African swine fever virus infection. Transbound Emerg Dis, 2018. 65(5): p. 1318–1328.

10. Schulz, K., et al., African Swine Fever in Saxony-Disease Dynamics. Viruses, 2024. 16(12).

11. Schäfer, A., et al., Adaptive Cellular Immunity against African Swine Fever Virus Infections. Pathogens, 2022. 11(2).

12. Mittelholzer, C., et al., Analysis of classical swine fever virus replication kinetics allows differentiation of highly virulent from avirulent strains. Veterinary microbiology, 2000. 74(4): p. 293–308.

13. Pietschmann, J., et al., Course and transmission characteristics of oral low-dose infection of domestic pigs and European wild boar with a Caucasian African swine fever virus isolate. Arch Virol, 2015. 160(7): p. 1657–67.

14. Rosenberg-Hasson, Y., et al., Effects of serum and plasma matrices on multiplex immunoassays. Immunol Res, 2014. 58(2-3): p. 224–33.

15. Sanchez-Cordon, P.J., et al., Absence of Long-Term Protection in Domestic Pigs Immunized with Attenuated African Swine Fever Virus Isolate OURT88/3 or BeninDeltaMGF Correlates with Increased Levels of Regulatory T Cells and Interleukin-10. J Virol, 2020. 94(14).

16. Netherton, C.L., et al., Identification and Immunogenicity of African Swine Fever Virus Antigens. Front Immunol, 2019. 10: p. 1318.

17. Tng, P.Y.L., et al., Capsid-Specific Antibody Responses of Domestic Pigs Immunized with Low-Virulent African Swine Fever Virus. Vaccines (Basel), 2023. 11(10).

18. Goatley, L.C., et al., Cellular and Humoral Immune Responses after Immunisation with Low Virulent African Swine Fever Virus in the Large White Inbred Babraham Line and Outbred Domestic Pigs. Viruses, 2022. 14(7).

19. Schwartz, J.C., et al., The major histocompatibility complex homozygous inbred Babraham pig as a resource for veterinary and translational medicine. HLA, 2018. 92(1): p. 40–3.

20. Schäfer, A., et al., T-cell responses in domestic pigs and wild boar upon infection with the moderately virulent African swine fever virus strain ‘Estonia2014’. Transbound Emerg Dis, 2021. 68(5): p. 2733–2749.

21. Radulovic, E., et al., The baseline immunological and hygienic status of pigs impact disease severity of African swine fever. PLoS Pathog, 2022. 18(8): p. e1010522.

22. Mellors, J., et al., Viral Evasion of the Complement System and Its Importance for Vaccines and Therapeutics. Front Immunol, 2020. 11: p. 1450.

23. Des Prez, R.M., et al., Function of the classical and alternate pathways of human complement in serum treated with ethylene glycol tetraacetic acid and MgCl2-ethylene glycol tetraacetic acid. Infect Immun, 1975. 11(6): p. 1235–43.

24. Boniotto, M., et al., Polymorphisms in the MBL2 promoter correlated with risk of HIV-1 vertical transmission and AIDS progression. Genes Immun, 2000. 1(5): p. 346–8.

25. Chong, W.P., et al., Mannose-binding lectin in chronic hepatitis B virus infection. Hepatology, 2005. 42(5): p. 1037–45.

26. Alves Pedroso, M.L., et al., Mannan-binding lectin MBL2 gene polymorphism in chronic hepatitis C: association with the severity of liver fibrosis and response to interferon therapy. Clin Exp Immunol, 2008. 152(2): p. 258–64.

27. Ip, W.K., et al., Mannose-binding lectin in severe acute respiratory syndrome coronavirus infection. J Infect Dis, 2005. 191(10): p. 1697–704.

28. Kwakkel-van Erp, J.M., et al., Mannose-binding lectin deficiency linked to cytomegalovirus (CMV) reactivation and survival in lung transplantation. Clin Exp Immunol, 2011. 165(3): p. 410–6.

29. Michelow, I.C., et al., High-dose mannose-binding lectin therapy for Ebola virus infection. J Infect Dis, 2011. 203(2): p. 175–9.

30. Tian, J., et al., Adenovirus activates complement by distinctly different mechanisms in vitro and in vivo: indirect complement activation by virions in vivo. J Virol, 2009. 83(11): p. 5648–58.

31. Gupta, P. and A.S. Tripathy, Alternative pathway of complement activation has a beneficial role against Chandipura virus infection. Med Microbiol Immunol, 2020. 209(2): p. 109–124.

32. Avirutnan, P., et al., Complement-mediated neutralization of dengue virus requires mannose-binding lectin. mBio, 2011. 2(6).

33. Fuchs, A., et al., Direct complement restriction of flavivirus infection requires glycan recognition by mannose-binding lectin. Cell Host Microbe, 2010. 8(2): p. 186–95.

34. Mason, C.P. and A.W. Tarr, Human lectins and their roles in viral infections. Molecules, 2015. 20(2): p. 2229–71.

